# Population-based analysis of ocular *Chlamydia trachomatis* in trachoma-endemic West African communities identifies genomic markers of disease severity

**DOI:** 10.1101/205336

**Authors:** AR Last, H Pickering, Ch Roberts, F Coll, J Phelan, SE Burr, E Cassama, M Nabicassa, HMB Seth-Smith, J Hadfield, LT Cutcliffe, IN Clarke, DCW Mabey, RL Bailey, TG Clark, NR Thomson, MJ Holland

## Abstract

*Chlamydia trachomatis* (*Ct*) is the most common infectious cause of blindness and bacterial sexually transmitted infection worldwide. Using *Ct* whole genome sequences obtained directly from conjunctival swabs, we studied *Ct* genomic diversity and associations between *Ct* genetic polymorphisms with ocular localization and disease severity in a treatment-naïve trachoma-endemic population in Guinea Bissau, West Africa. All sequences fall within the T2 ocular clade phylogenetically. This is consistent with the presence of the characteristic deletion in *trpA* resulting in a truncated non-functional protein and the ocular tyrosine repeat regions present in *tarP* associated with ocular tissue localization. We have identified twenty-one *Ct* non-synonymous single nucleotide polymorphisms (SNPs) associated with ocular localization, including SNPs within *pmpD* (OR=4.07, *p*=0.001*) and *tarP* (OR=0.34, *p*=0.009*). Eight SNPs associated with disease severity were found in *yjfH (rlmB)* (OR=0.13, *p*=0.037*), *CTA0273* (OR=0.12, *p*=0.027*), *trmD* (OR=0.12, *p*=0.032*), *CTA0744* (OR=0.12, *p*=0.041*), *glgA* (OR=0.10, *p*=0.026*), *alaS* (OR=0.10, *p*=0.032*), *pmpE* (OR=0.08, *p*=0.001*) and the intergenic region *CTA0744-CTA0745* (OR=0.13, *p*=0.043*). This study demonstrates the extent of genomic diversity within a naturally circulating population of ocular *Ct*, and the first to describe novel genomic associations with disease severity. These findings direct investigation of host-pathogen interactions that may be important in ocular *Ct* pathogenesis and disease transmission.

## INTRODUCTION

The obligate intracellular bacterium *Chlamydia trachomatis (Ct)* is the leading infectious cause of blindness (trachoma) and the most common sexually transmitted bacterial infection^1,2^.

*Ct* strains are differentiated into biovars based on pathobiological characteristics and serovars based on serological reactivity for the major outer membrane protein (MOMP) encoded by *ompA*^3^. Serovars largely differentiate biological groups associated with trachoma (A-C), sexually transmitted disease (D-K) and lymphogranuloma venereum (LGV) (L1-3). Despite diverse biological phenotypes, *Ct* strains share near complete genomic synteny and gene content^4^, suggesting that minor genetic changes influence pathogen-host and tissue-specific infection characteristics^5–7^. All published African ocular *Ct* genomes are situated on the ocular branch within the T2 clade of non-LGV urogenital isolates^4^. Currently there are only 31 published ocular *Ct* genome sequences^4,9–12^. In particular there appears to be limited genomic diversity between published *Ct* genomes from Gambian and Tanzanian populations^8^.

The pathogenesis of chlamydial infection begins with epithelial inflammation and may progress to chronic immuno-fibrogenic processes leading to blindness and infertility, though many *Ct* infections do not result in sequelae^13,14^. Strain-specific differences related to clinical presentation have been investigated in trachoma^8,15,16^. These studies examined a small number of ocular *Ct* isolates from the major trachoma serotypes and found a small subset of genes in addition to *ompA* that were associated with differences in *in vitro* growth rate, burst size, plaque morphology, interferon-γ sensitivity and most importantly, intensity of infection and clinical disease severity in non-human primates (NHPs), suggesting that genetic polymorphisms in *Ct* may contribute to the observed variability in severity of trachoma in endemic communities^8^.

The obligate intracellular development of *Ct* has presented significant technical barriers to basic research into chlamydial biology. Only recently has genetic manipulation of the chlamydial plasmid been possible, allowing *in vitro* transformation and modification studies, though this remains technically challenging, necessitating alternative approaches^17,18^.

Whole genome sequencing (WGS) has recently been used to identify regions of likely recombination in recent clinical isolates, demonstrating that WGS analysis may be an effective approach for the discovery of loci associated with clinical presentation^6^. Additionally, a number of putative virulence factors have been identified through WGS analysis and subsequent *in vitro* and animal studies^5,19–30^. However there are currently no published population-based studies of *Ct* using WGS with corresponding detailed clinical data, making it difficult to relate genetic changes to functional relevance and virulence factors *in vivo*.

There is an increasing pool of *Ct* genomic data, largely from archived samples following cell culture and more recently directly from clinical samples^31^. WGS data obtained directly from clinical samples can be preferable to using WGS data obtained from cell cultured *Ct*, since repeated passage of *Ct* results in mutations that are not observed *in vivo*^32–34^.

*Ct* bacterial load is associated with disease severity, particularly conjunctival inflammation, in active (infective) trachoma^35^. Conjunctival inflammation has previously been shown to be a marker of severe disease and plays an important role in the pathogenesis of scarring trachoma^36–38^. In this study we used principal component analysis (PCA) to reduce the dimensions of clinical grade of inflammation (defined using the P score from the FPC trachoma grading system^39^) and *Ct* bacterial load to a single metric to define an *in vivo* conjunctival phenotype in active (infective) trachoma. PCA is a recognised dimension reduction technique used to combine multiple correlated traits into their uncorrelated principal components (PC)^40–42^, allowing us to examine the relationship between *Ct* genotype and disease severity. These data currently represent the largest collection of ocular *Ct* sequences from a single population from the trachoma-endemic region of the Bijagós Archipelago of Guinea Bissau, and provide a unique opportunity to gain insight into ocular *Ct* pathogenesis in humans.

## RESULTS

Conjunctival swabs collected during a cross-sectional population-based trachoma survey on the Bijagós Archipelago yielded 220 ocular *Ct* infections detected by *Ct* plasmid-based droplet digital PCR (ddPCR). Of the 220 *Ct* infections detected, 184 were quantifiable using *Ct* genome-based ddPCR.

We obtained WGS data from 126 using cell culture (*n=8*) or direct sequencing from swabs with SureSelect^XT^ target enrichment (*n=118*), representing the largest cross-sectional collection of ocular *Ct* WGS. Eighty-one of these sequences were subsequently included in the phylogenetic and diversity analyses and 71 were retained in the final genome-wide association (tissue localization (derived from the anatomical site of sample collection) and disease severity) analyses. The quality filtering process is illustrated in *Figure 1* and detailed in Methods. Briefly, we used standard GATK SNP-calling algorithms where >10× mean depth of coverage is defined as a threshold value and performs well in variant calling, is highly sensitive and has a false positive rate of <0.05%^43,44^.

**Figure 1.**
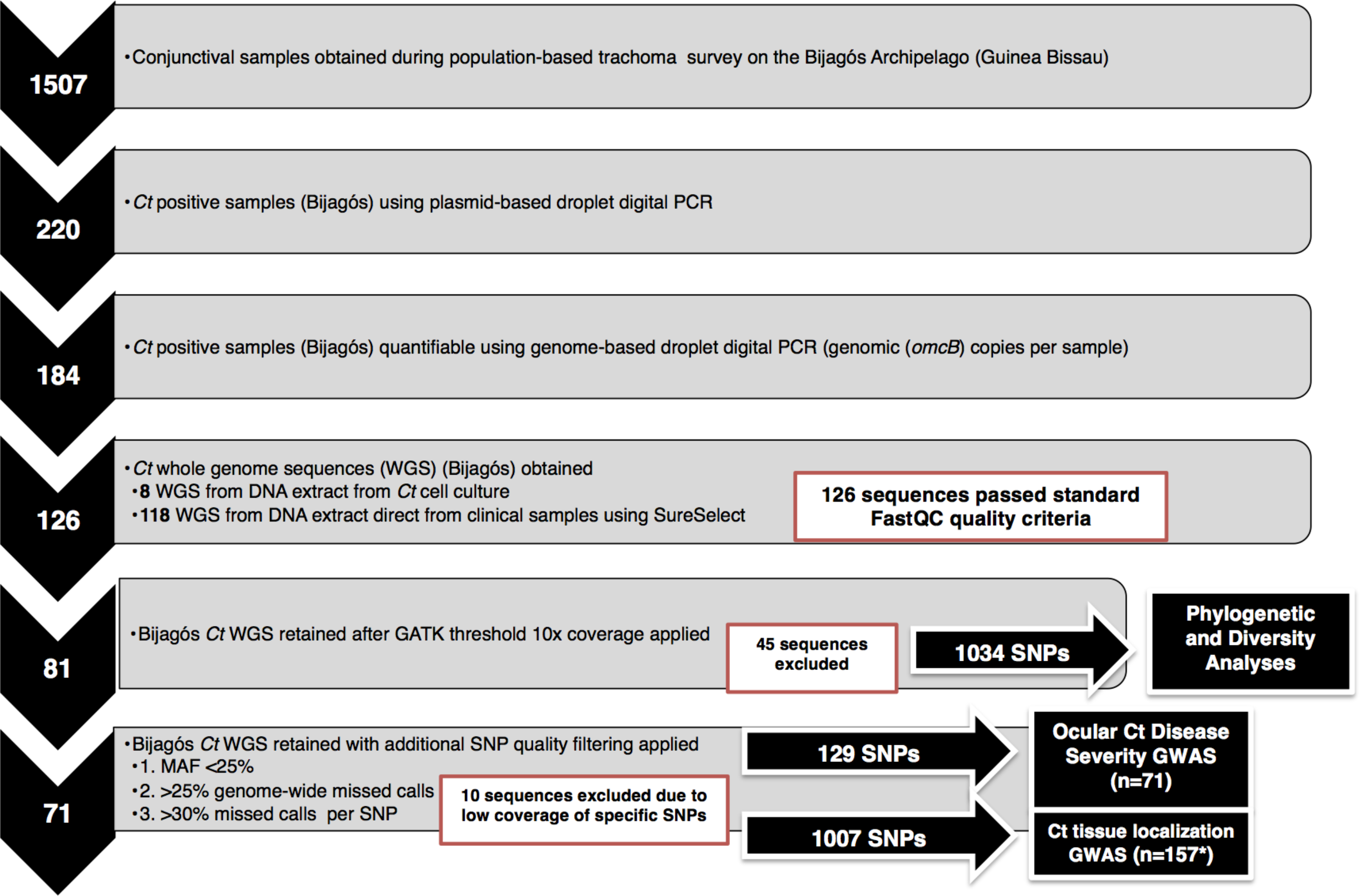
Whole Genome Sequence (WGS) quality filtering processes and threshold criteria for inclusion in analyses *Ct* DNA detected using droplet digital PCR ^82^. WGS data were obtained using SureSelect target enrichment^31^ (or chlamydial cell culture) and Illumina paired end sequencing. FastQC^90^ was used to assess basic WGS quality. SNP alleles were called against reference strain *Ct A/HAR-13* using an alternative coverage- based approach where a missing call was assigned to a site if the total coverage was less than 20x depth or where one of the four nucleotides accounted for at least 80% total coverage^95^. There was a clear relationship between the mean depth of coverage and genome-wide proportion of missing calls, therefore only sequences with greater than 10x mean depth of coverage over the whole genome were retained using the GATK Best Practice threshold^93,94^. Heterozygous calls were removed and SNPs with a minor allele frequency (MAF) of less than 25% were removed. Samples with greater than 25% genome-wide missing data and 30% missing data per SNP were excluded from the analysis. WGS sequence quality is shown in detail in *Supplementary Information S12*. *n=157 including the 71 Bijagós sequences in addition to 48 Rombo sequences and 38 reference sequences.

A total of 1034 unique SNP sites were identified within the 126 Bijagós *Ct* genomes relative to the reference strain *Ct A/HAR-13*. Following application of further threshold criteria based on minor allele frequency (MAF) and genome-wide missing data thresholds, we retained only high quality genomic data in the final association analyses (129 SNPs from 71 sequences). There were no significant differences between the 71 retained and the 55 excluded sequences with respect to demographic characteristics, bacterial load, disease severity scores or geographical location (*Table 1*). Clinical and demographic details of the survey participants in whom we did not identify *Ct* infection have been published previously^45^. Of the ten SNPs initially identified within the *Ct* plasmid sequences, none fulfilled the quality filtering criteria and were not retained for the genome-wide association analyses.

**Table 1.** Characteristics of ocular *Chlamydia trachomatis* sequences included in the disease severity association analysis Sequences (n=55) were excluded from the association analysis if there was a) <10x coverage b)* >25% missing reads genome-wide and c) >25% missing (N) calls at the single nucleotide polymorphism (SNP) locus. Coverage and missing data were correlated and resulted in exclusion of the same samples irrespective of criteria chosen. 71 sequences were retained in the final disease severity analysis. **Ocular *C. trachomatis* load = *omcB* (*C. trachomatis* genome) copies per conjunctival swab measured using droplet digital PCR. *** P score = Conjunctival inflammation score (0-3) using the modified FPC (Follicles, Papillary Hypertrophy, Conjunctival Scarring) grading system for trachoma^39^.

| Sequence ID | Sample ID | Average Depth of Coverage | % Missing Reads* | Gender | Age (years) | Island of Origin | Village Code | Ocular Load** | P Score* |
| --- | --- | --- | --- | --- | --- | --- | --- | --- | --- |
| 11152_3_1 | 14344 | 764 | 0.35% | M | 4 | Canhabaque | 33 | 202632 | 1 |
| 11152_3_10 | 17347 | 121 | 0.21% | M | 5 | Bubaque | 17 | 69093 | 2 |
| 11152_3_11 | 4422 | 19 | 19.95% | F | 2 | Bubaque | 12 | 68782 | 2 |
| 11152_3_12 | 11231 | 68 | 2.24% | M | 0 | Soga | 43 | 64036 | 1 |
| 11152_3_13 | 15631 | 21 | 14.93% | F | 2 | Canhabaque | 33 | 55749 | 3 |
| 11152_3_14 | 6105 | 1664 | 0.05% | F | 1 | Bubaque | 14 | 55202 | 3 |
| 11152_3_15 | 12628 | 191 | 0.10% | F | 12 | Canhabaque | 29 | 54651 | 2 |
| 11152_3_16 | 7524 | 2065 | 0.14% | M | 10 | Canhabaque | 35 | 54539 | 2 |
| 11152_3_17 | 5016 | 61 | 0.44% | F | 1 | Bubaque | 15 | 46510 | 2 |
| 11152_3_18 | 1485 | 44 | 1.21% | F | 4 | Canhabaque | 27 | 45929 | 1 |
| 11152_3_19 | 15554 | 825 | 0.06% | F | 1 | Canhabaque | 33 | 44052 | 2 |
| 11152_3_20 | 6094 | 3070 | 0.00% | F | 3 | Bubaque | 14 | 42917 | 2 |
| 11152_3_22 | 5082 | 51 | 0.81% | M | 6 | Bubaque | 15 | 42427 | 1 |
| 11152_3_23 | 12969 | 3643 | 1.81% | F | 3 | Canhabaque | 29 | 41308 | 3 |
| 11152_3_25 | 8140 | 246 | 0.36% | M | 13 | Bubaque | 20 | 39816 | 2 |
| 11152_3_26 | 6083 | 2746 | 0.00% | F | 23 | Bubaque | 14 | 38771 | 3 |
| 11152_3_27 | 16621 | 1664 | 0.00% | M | 3 | Canhabaque | 37 | 33514 | 3 |
| 11152_3_28 | 16852 | 143 | 0.16% | M | 5 | Canhabaque | 38 | 31228 | 2 |
| 11152_3_29 | 16588 | 53 | 0.81% | M | 6 | Canhabaque | 37 | 29991 | 1 |
| 11152_3_3 | 4180 | 51 | 0.92% | M | 2 | Bubaque | 12 | 140693 | 2 |
| 11152_3_30 | 7612 | 107 | 0.44% | F | 3 | Canhabaque | 35 | 28528 | 2 |
| 11152_3_31 | 6985 | 177 | 0.10% | M | 6 | Bubaque | 17 | 27924 | 2 |
| 11152_3_32 | 4411 | 24 | 9.68% | F | 1 | Bubaque | 12 | 27584 | 2 |
| 11152_3_33 | 4257 | 381 | 0.06% | M | 0 | Bubaque | 12 | 24033 | 3 |
| 11152_3_34 | 4400 | 48 | 0.98% | M | 6 | Bubaque | 12 | 23435 | 2 |
| 11152_3_35 | 15180 | 571 | 0.35% | F | 7 | Canhabaque | 33 | 23254 | 0 |
| 11152_3_36 | 13596 | 496 | 0.06% | M | 18 | Canhabaque | 23 | 22098 | 3 |
| 11152_3_37 | 1672 | 20 | 18.42% | M | 6 | Canhabaque | 25 | 21630 | 3 |
| 11152_3_38 | 5181 | 81 | 0.32% | M | 4 | Bubaque | 15 | 21339 | 2 |
| 11152_3_39 | 15532 | 243 | 0.08% | F | 25 | Canhabaque | 33 | 21174 | 2 |
| 11152_3_4 | 8074 | 150 | 0.13% | M | 4 | Bubaque | 18 | 131175 | 2 |
| 11152_3_40 | 16984 | 145 | 0.19% | M | 4 | Canhabaque | 21 | 20113 | 1 |
| 11152_3_41 | 1881 | 37 | 2.71% | F | 1 | Canhabaque | 32 | 15963 | 2 |
| 11152_3_42 | 10032 | 101 | 0.16% | M | 2 | Soga | 42 | 15706 | 1 |
| 11152_3_43 | 8492 | 70 | 2.60% | M | 1 | Rubane | 45 | 15582 | 2 |
| 11152_3_44 | 13585 | 31 | 4.97% | M | 23 | Canhabaque | 23 | 15417 | 3 |
| 11152_3_48 | 7535 | 61 | 0.84% | M | 18 | Canhabaque | 35 | 13439 | 3 |
| 11152_3_5 | 7095 | 235 | 0.44% | F | 4 | Bubaque | 17 | 105453 | 3 |
| 11152_3_50 | 6028 | 46 | 1.24% | F | 4 | Bubaque | 14 | 12961 | 2 |
| 11152_3_52 | 10021 | 20 | 16.15% | F | 6 | Soga | 42 | 11840 | 1 |
| 11152_3_55 | 12650 | 59 | 0.54% | M | 6 | Canhabaque | 29 | 9001 | 2 |
| 11152_3_57 | 8965 | 21 | 16.60% | M | 27 | Soga | 43 | 7336 | 1 |
| 11152_3_58 | 5104 | 33 | 3.68% | M | 2 | Bubaque | 15 | 7203 | 2 |
| 11152_3_6 | 16599 | 52 | 0.73% | M | 9 | Canhabaque | 37 | 96333 | 2 |
| 11152_3_62 | 7062 | 22 | 13.41% | F | 4 | Bubaque | 17 | 6986 | 3 |
| 11152_3_63 | 8778 | 17 | 25.47% | F | 11 | Rubane | 46 | 6760 | 3 |
| 11152_3_66 | 1892 | 45 | 1.25% | F | 2 | Canhabaque | 32 | 6374 | 1 |
| 11152_3_7 | 10747 | 581 | 1.82% | F | 3 | Soga | 44 | 82916 | 2 |
| 11152_3_70 | 13189 | 25 | 8.87% | F | 3 | Canhabaque | 24 | 4703 | 1 |
| 11152_3_74 | 15499 | 24 | 10.49% | M | 5 | Canhabaque | 33 | 4226 | 1 |
| 11152_3_76 | 726 | 417 | 0.06% | F | 3 | Canhabaque | 26 | 3753 | 0 |
| 11152_3_77 | 7579 | 105 | 0.52% | F | 5 | Canhabaque | 35 | 3468 | 1 |
| 11152_3_78 | 12089 | 16 | 27.78% | F | 13 | Canhabaque | 47 | 3203 | 2 |
| 11152_3_8 | 6996 | 38 | 2.03% | M | 3 | Bubaque | 17 | 82614 | 1 |
| 11152_3_88 | 748 | 163 | 0.10% | F | 2 | Canhabaque | 26 | 1636 | 0 |
| 11152_3_9 | 10967 | 20 | 17.52% | F | 2 | Soga | 44 | 81124 | 3 |
| 11152_3_92 | 1463 | 73 | 0.30% | F | 42 | Canhabaque | 27 | 1273 | 2 |
| 13108_1_14 | 24519 | 51 | 2.81% | M | 2 | Rubane | 45 | 29040 | 3 |
| 13108_1_15 | 6941 | 33 | 1.81% | M | 36 | Bubaque | 17 | 13155 | 1 |
| 13108_1_7 | 25124 | 27 | 5.27% | M | 4 | Canhabaque | 22 | 21750 | 3 |
| 13108_1_9 | 22154 | 18 | 20.56% | F | 5 | Soga | 43 | 14349 | 1 |
| 8422_8_49 | 2353 | 39 | 5.70% | M | 11 | Canhabaque | 35 | 96889 | 2 |
| 8422_8_50 | 2366 | 82 | 1.08% | M | 1 | Canhabaque | 35 | 289778 | 2 |
| 9471_4_86 | 12980 | 287 | 1.90% | M | 4 | Canhabaque | 29 | 85456 | 1 |
| 9471_4_87 | 15367 | 215 | 0.46% | M | 1 | Canhabaque | 33 | 99064 | 1 |
| 9471_4_88 | 15543 | 192 | 0.11% | F | 23 | Canhabaque | 33 | 49125 | 1 |
| 9471_4_89 | 1870 | 119 | 0.14% | M | 3 | Canhabaque | 32 | 158548 | 3 |
| 9471_4_90 | 2145 | 111 | 0.11% | M | 15 | Canhabaque | 32 | 140297 | 2 |
| 9471_4_91 | 4158 | 94 | 0.14% | M | 4 | Bubaque | 12 | 63654 | 1 |
| 9471_4_92 | 4169 | 85 | 0.13% | F | 3 | Bubaque | 12 | 274835 | 2 |
| 9471_4_93 | 7590 | 242 | 0.51% | F | 1 | Canhabaque | 35 | 128025 | 3 |

### Ocular C. trachomatis Phylogeny and Diversity

For the phylogeny and diversity analyses 81 Bijagós *Ct* sequences were included on the basis of quality filtering criteria described in detail in *Figure 1*. SNP-based phylogenetic trees constructed using all 1034 SNPs for sequences above 10× coverage (*n=81*), with 54 published *Ct* reference genomes, are shown in *Figure 2*.

**Figure 2.**
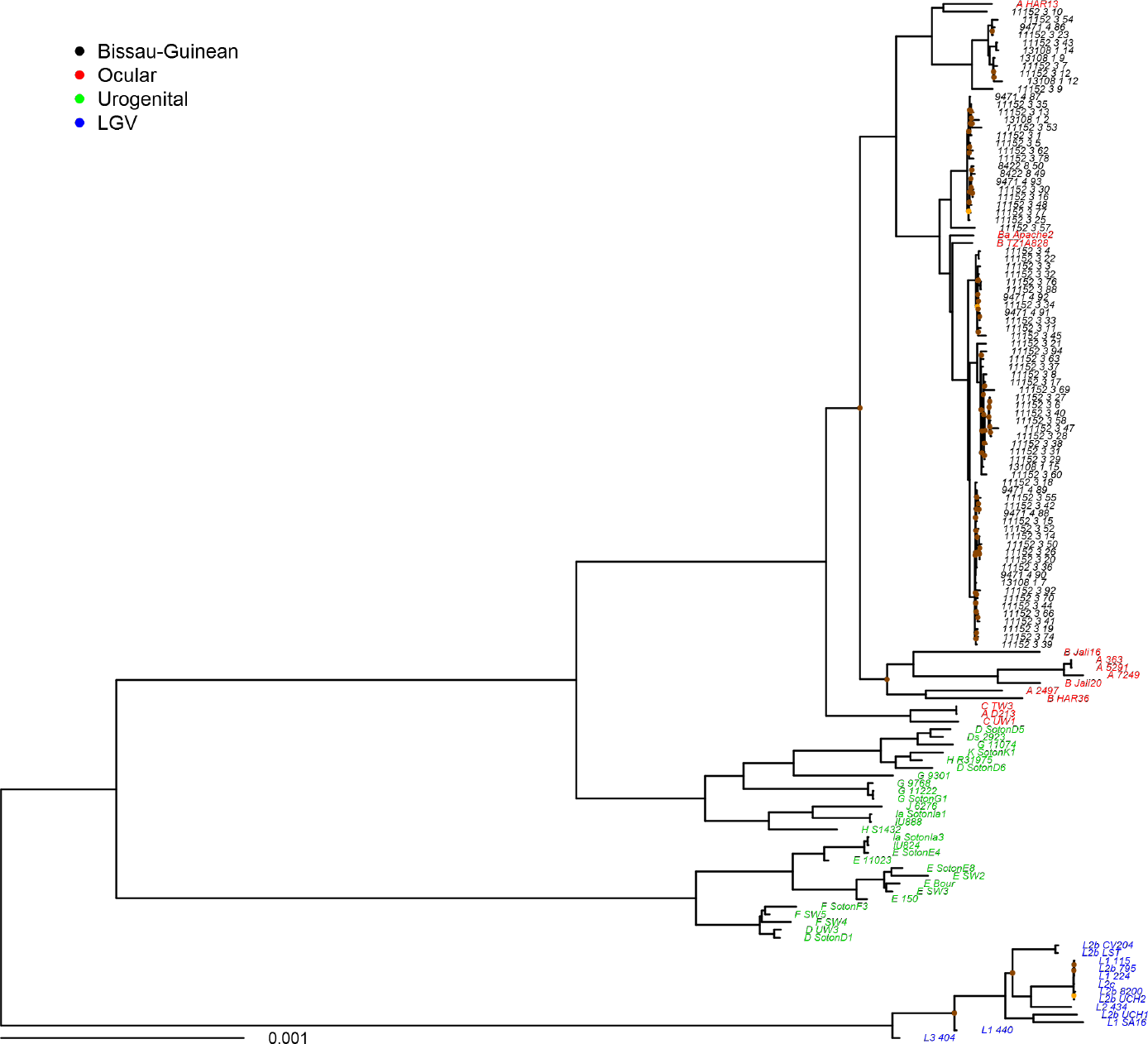
Maximum likelihood reconstruction of whole-genome phylogeny of ocular *Chlamydia trachomatis* sequences from the Bijagós Archipelago (Guinea Bissau) Maximum likelihood reconstruction of the whole-genome phylogeny of 81 *Ct* sequences from the Bijagós Islands and 54 *Ct* reference strains. Bijagós *Ct* sequences (n=81) were mapped to *Ct A/HAR-13* using SAMtools^92^. SNPs were called as described by Harris et al.^4^ Phylogenies were computed with RAxML^96^ from a variable sites alignment using a GTR+gamma model and are midpoint rooted. The scale bar indicates evolutionary distance. Bijagós *Ct* sequences in this study are coloured BLACK and reference strains are coloured by tissue localization (RED=Ocular, GREEN=Urogenital, BLUE=LGV). Branches are supported by > 90% of 1000 bootstrap replicates. Branches supported by 80-90% (ORANGE) and < 80% (BROWN) bootstrap replicates are indicated.

The Bijagós sequences are situated within the T2 ocular monophyletic lineage with all other ocular *Ct* sequences^46^ except those described by Andersson *et al*.^10^. However, our population-based collection of ocular *Ct* sequences has much greater diversity at whole genome resolution than previously demonstrated in African trachoma isolates^4,8^. We used a pairwise diversity (π) metric to compare two populations of ocular *Ct* from regions with similar trachoma endemicity and studies with similar design, sample size and available epidemiological metadata. These data show much greater genomic diversity in the Bijagós ocular *Ct* sequences (π=0.07167) compared to the Tanzanian (Rombo) ocular *Ct* sequences (π =0.00047).

By *ompA* genotyping, 73 of the Bijagós sequences are genotype A and eight are genotype B, supporting their classical ocular nature (*Supplementary Information S1*). The high resolution of WGS data obtained directly from clinical samples captures diversity that may be useful in strain classification, particularly as we found some evidence of clustering at village level, although the very small number of sequences per village means that it is not possible to provide accurate estimates of clustering in this study (*Figure 3*).

**Figure 3.**
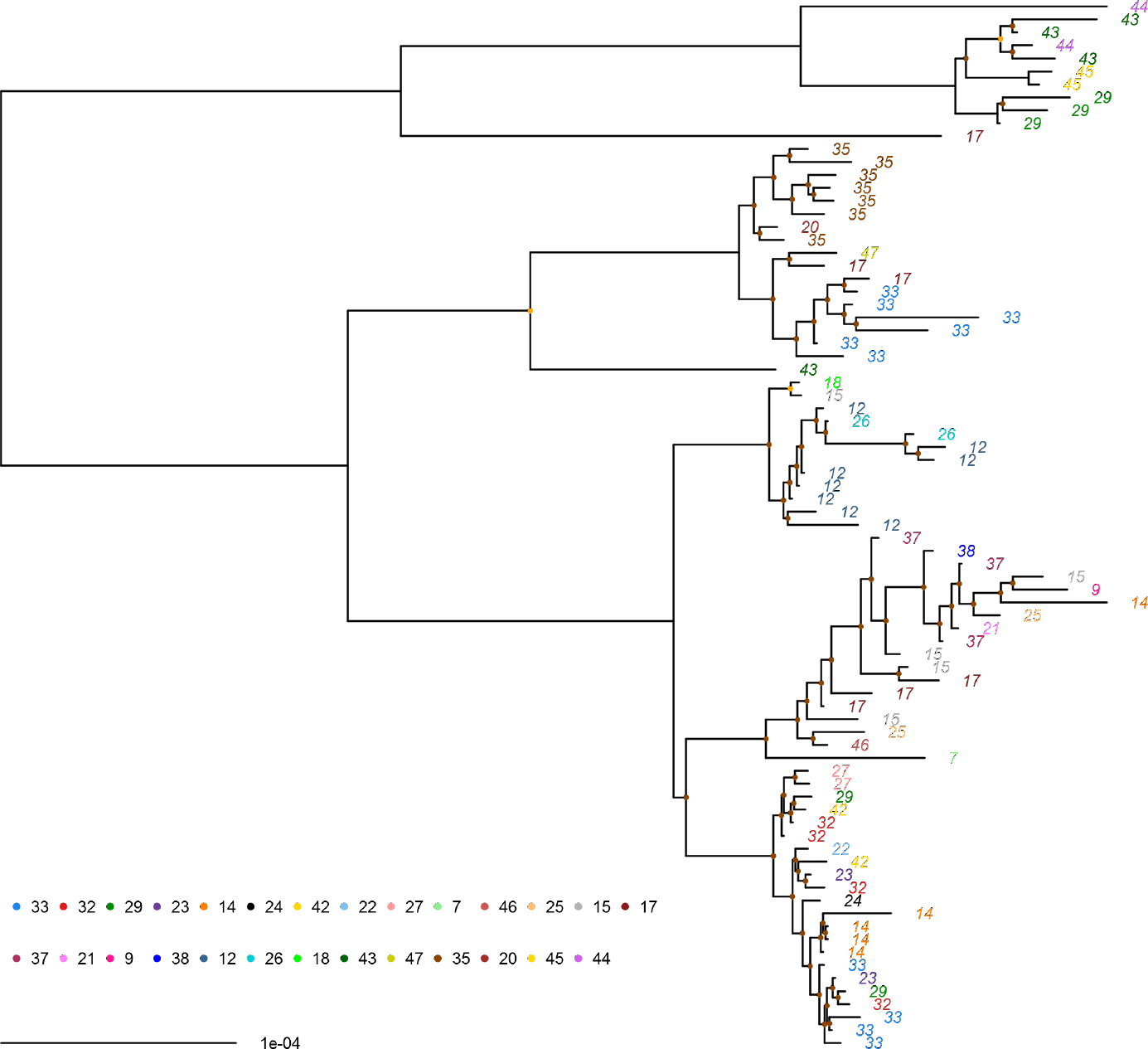
Maximum likelihood phylogenetic tree showing clustering of ocular *Chlamydia trachomatis* sequence types by village. RAxML maximum likelihood phylogenetic reconstruction including all ocular *Ct* sequences retained in the final disease severity association analysis after quality filtering (*n=71*). Ocular *Ct* sequences labelled by village (villages numbered and coloured). Midpoint-rooted and mapped to reference *Ct A/HAR-13*. Branches are supported by > 90% of 1000 bootstrap replicates. Branches supported by 80-90% (ORANGE) and < 80% (BROWN) bootstrap replicates are indicated.

Homoplasic SNPs and regions affected by recombination are shown in *Supplementary Information S2 (a)*. Removal of these regions of recombination identified using the pairwise homoplasy index had no effect on phylogenetic relationships. Additionally, a site-wise log likelihood plot demonstrated that there was no clear genomic region where there was significant lack of confidence in the tree construction due to recombination (*Supplementary Information S2 (b)*). Whether regions containing recombination were included or excluded, tree topology remained essentially identical, indicating that branching order is not affected by the removal of these regions.

### Genome-wide analysis of C. trachomatis localization

Candidate genes thought to be involved in or indicative of ocular localization or preference were examined to further characterize this population of ocular *Ct*. Polymorphisms and truncations in the tryptophan operon have previously been implicated in the inability of ocular *Ct* to infect and survive in the genital tract^5^. All sequences contained mutations in *trpA* resulting in truncation. The majority (80/81) were truncated at the previously characterised deletion at position 533^5^. Polymorphims in *trpB* and *trpR* were less common (*Supplementary Information S3*).

The variable domain structure of the translocated actin-recruiting phosphoprotein (*tarP*) has also been implicated in tropism^47^. Ocular strains possess more actin-binding domains (three or four) and fewer tyrosine repeat regions (between one and three). Urogenital strain *tarP* sequences have low copy numbers of both and LGV strain sequences have additional tyrosine repeat regions. In this study, all sequences contain the expected three tyrosine repeat regions and three or four actin-binding domains (*Supplementary Information S3*).

The nine virulence associated polymorphic membrane proteins (Pmp) are variably related to tissue preference with all encoding genes except *pmpA*, *pmpD* and *pmpE* clustering by tissue location^20^. In this population all phylogenies of the six tropism-clustering *pmps* show that all sequences cluster with other ocular sequences (*Supplementary Information S4*).

Permutation-based re-sampling methods, commonly used in GWAS analyses, were used to account for multiple comparisons^48–51^. 1007 SNPs were tested in 157 *Ct* sequences (*Figure 1*) for association with ocular localization (defined by anatomical site of sample collection), comparing eight ocular, 17 urogenital and 13 LGV strains (*Figure 4(a)*). One hundred and five SNPs were significantly associated with ocular localization (*p*<0.05*) of which 21 were non-synonymous (details in *Table 2(a) and Supplementary Information S5*). These were within a number of genes known to be polymorphic, genes previously identified as tropism-associated (*CTA0156*, *CTA0498*/*tarP* and *CTA0743*/*pbpB*) and virulence factors (*CTA0498*/*tarP* and *CTA0884*/*pmpD*). No predicted protein localization was over-represented in the ocular localization-related SNPs (*p=0.6174*), however early and very-late expressed genes were over-represented (*p=0.0197*).

**Table 2.**
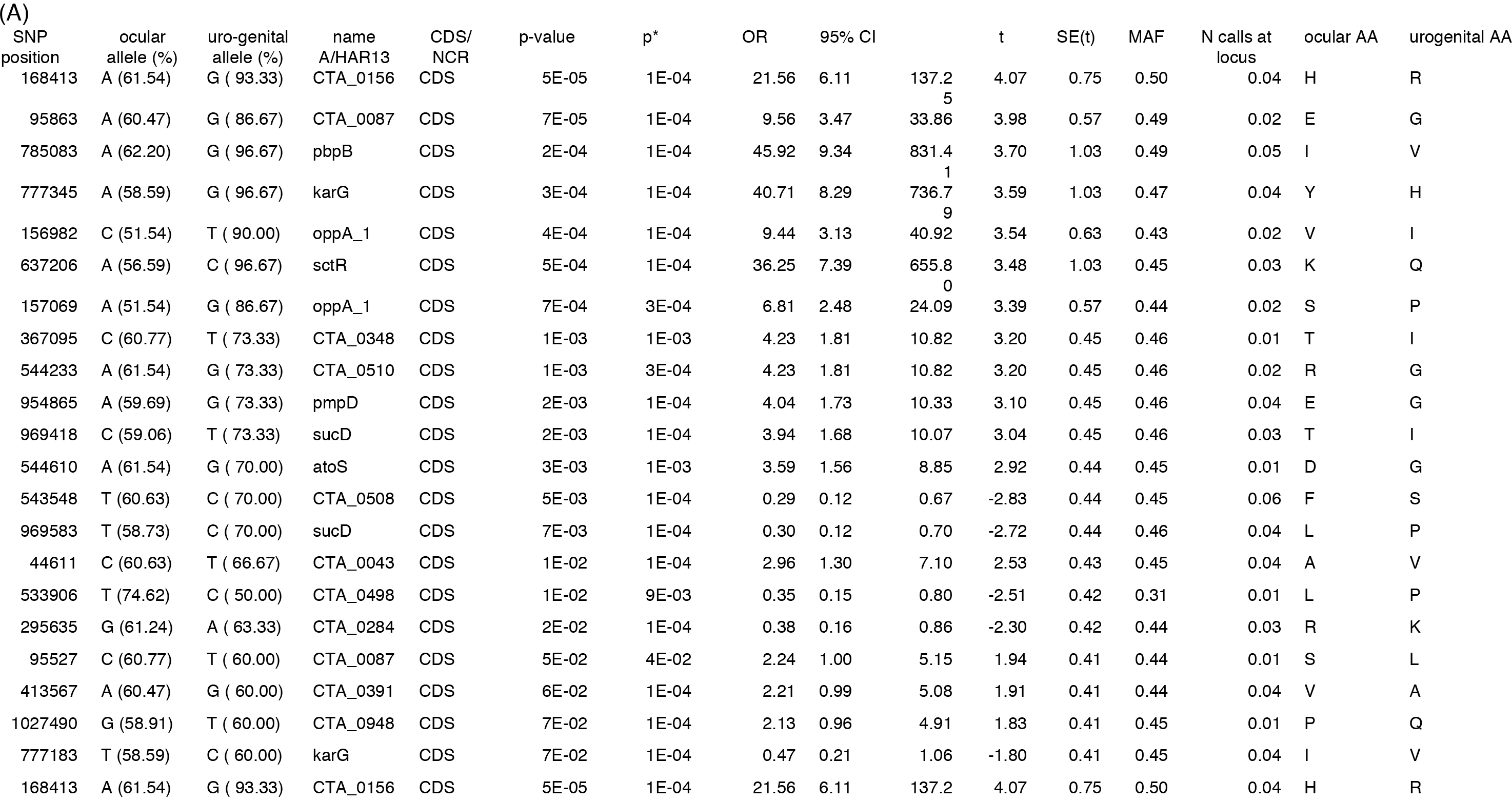
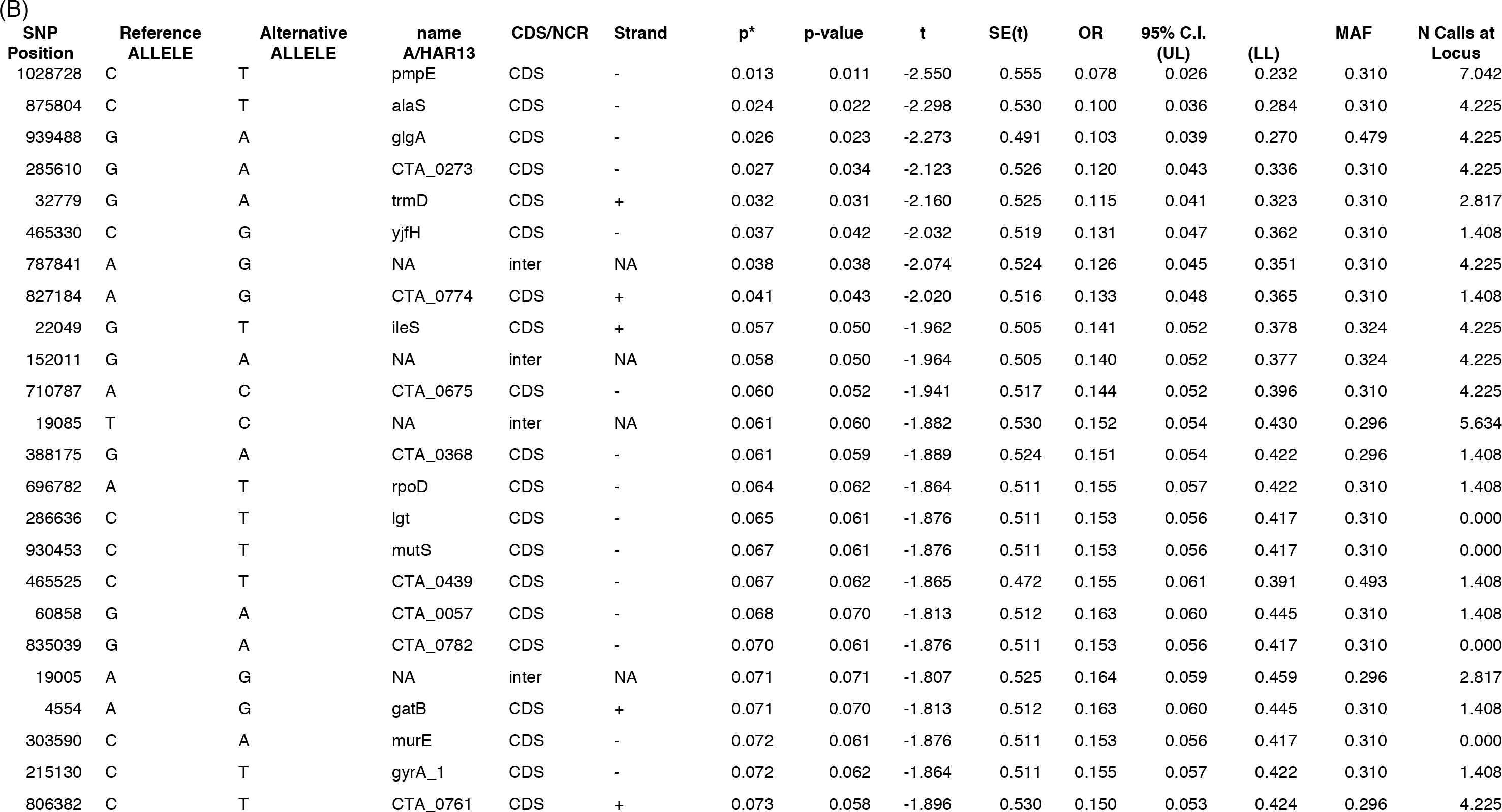
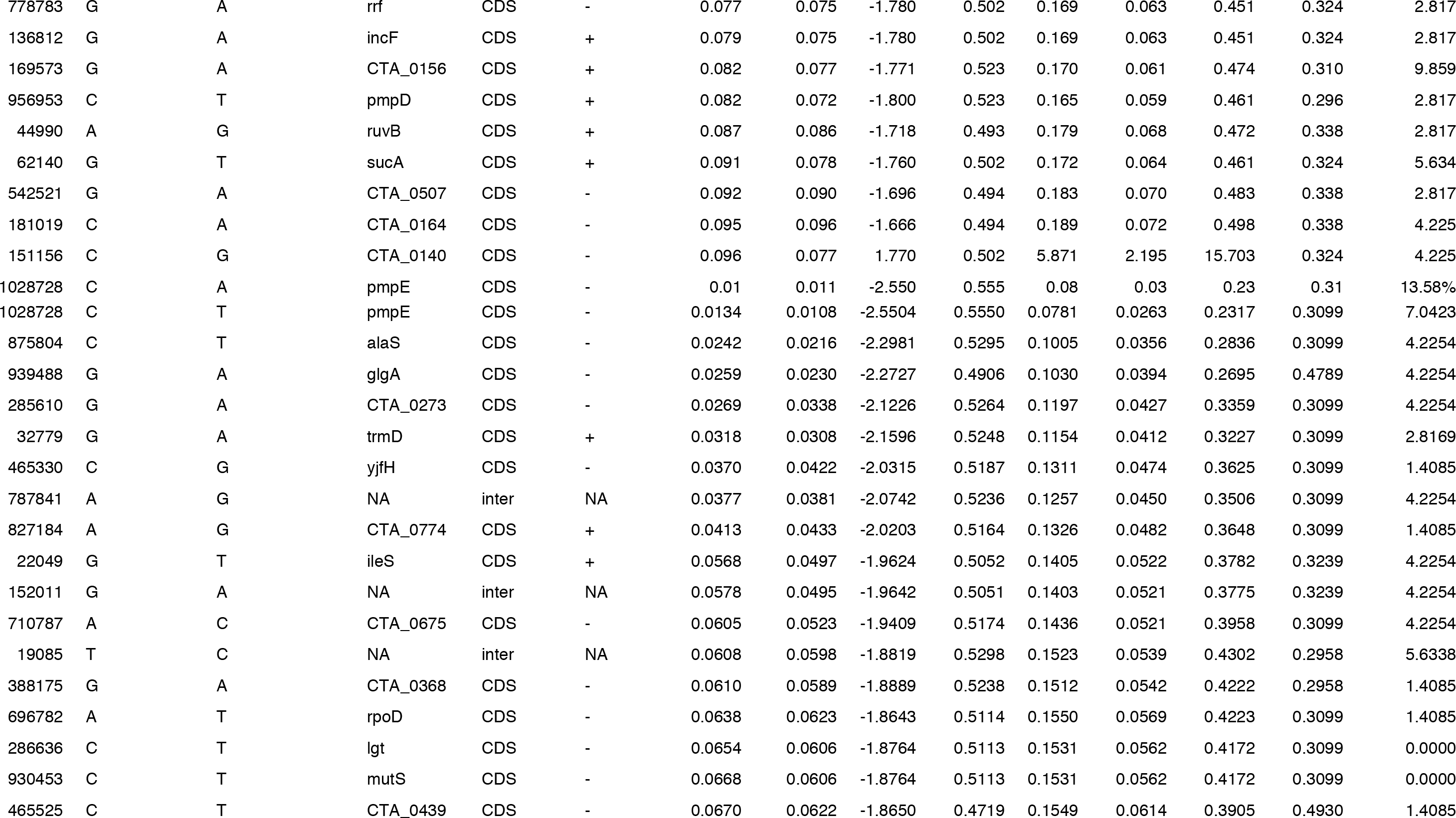
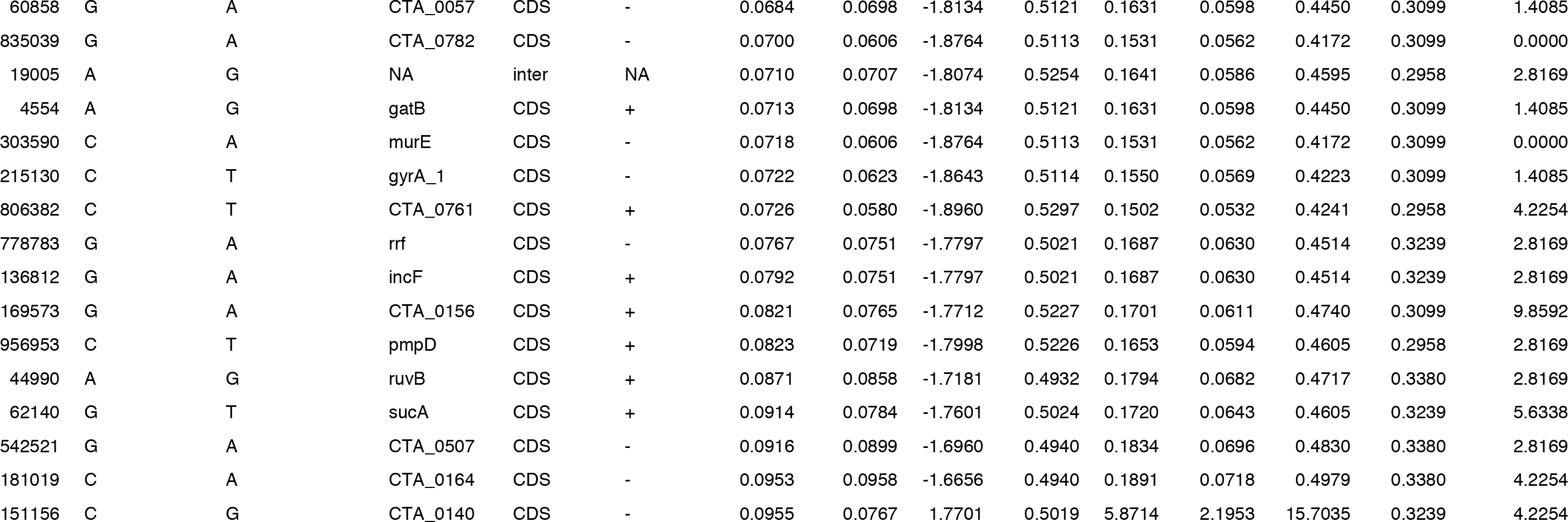
SNPs across the *Chlamydia trachomatis* genome identified using permutation-based genome-wide association analysis for (A) ocular localization (non-synonymous only) and (B) disease severity (a) Ocular localization-associated non-synonymous SNPs (*p-value* < 0.1). Position of the SNPs and name of the impacted are from the *Ct A/HAR13* (GenBank Accession Number NC_007429) genome. ‘Allele Percentage’ is the percentage of each group where the given allele was present. ‘CDS/NCR’ identifies whether the SNP was in a coding or non-coding region. ‘P*’ indicates p-values from 100,024 simulations indicating genome wide significance at *p*\*<0.05. ‘MAF’ is the minor allele frequency. ‘N Calls at Locus’ is the proportion of isolates which had no base called. ‘AA’ is the amino acid coded for. (b) Disease severity-associated SNPs (*p-value* < 0.1). Disease severity is defined by a composite in vivo conjunctival phenotype derived using principal component analysis using ocular C. trachomatis load and conjunctival inflammatory (P) score (using the modified FPC (Follicles, Papillary Hypertrophy, Conjunctival Scarring) trachoma grading system39). ‘Reference Allele’ indicates the reference allele on *Ct A/HAR-13* (GenBank Accession Number NC_007429). ‘CDS/NCR’ identifies whether the SNP was in a coding or non-coding region. ‘P*’=permuted p-value after 100,024 simulations indicating genome wide significance at *p*\*<0.05. ‘T’ is the t statistic; SE(T) is the Standard Error of the t statistic. OR is the adjusted Odds Ratio (derived from the t statistic). 95% C.I.=95% confidence interval of the OR. ‘MAF’ is the minor allele frequency. ‘N Calls at Locus’ is the proportion of isolates which had no base called.

**Figure 4.**
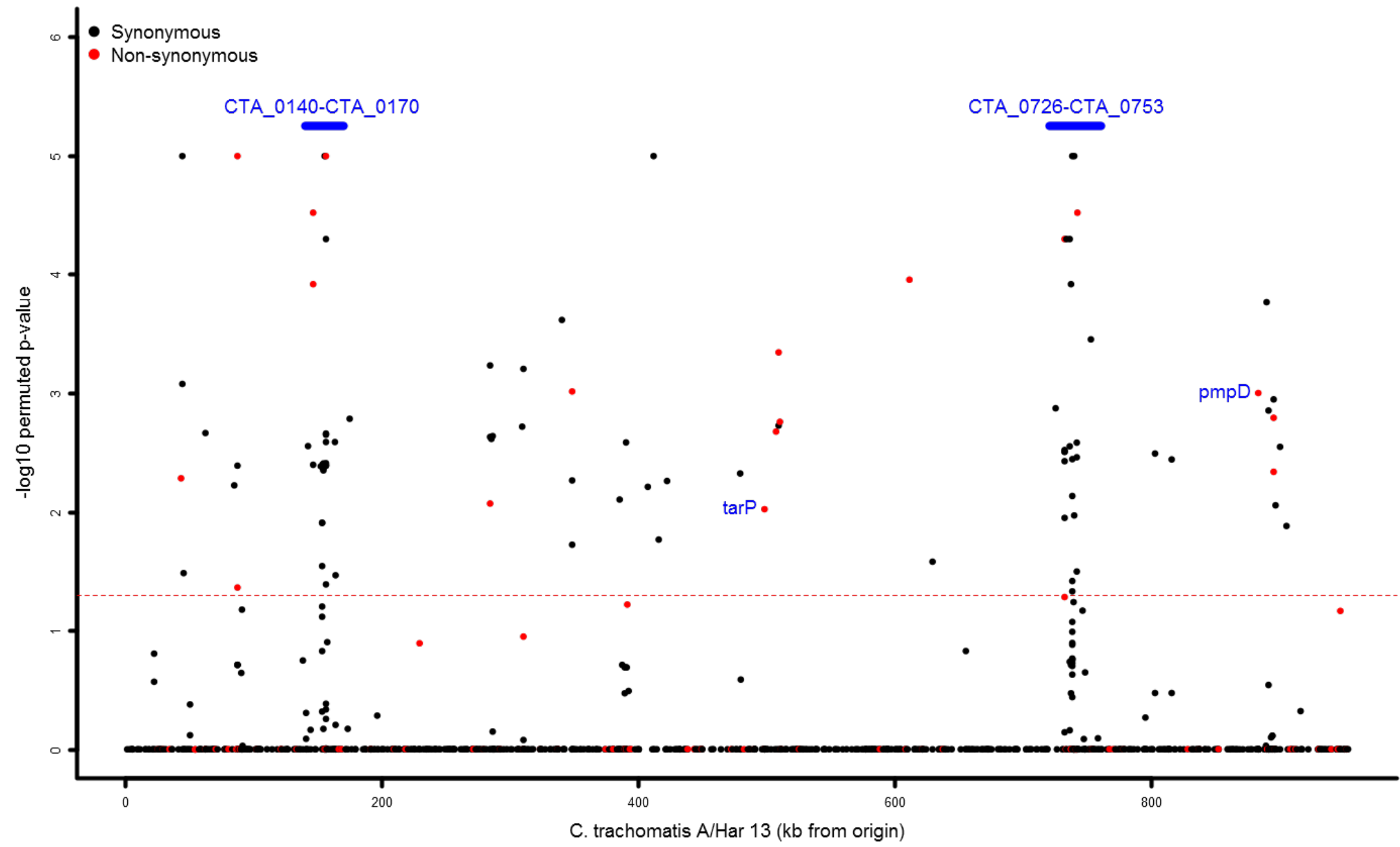
Single Nucleotide Polymorphisms on the *Chlamydia trachomatis* genome associated with (A) ocular localization and (B) disease severity at genome-wide significance. (A) Ocular localization-associated SNPs across the *C. trachomatis* genome. 1007 SNPs were identified in coding and non-coding regions and were included in permutation-based linear regression models in the *Ct* genome-wide association analysis. The threshold for genome-wide significance is indicated by the dashed line (*p*<0.05*). The y-axis shows the −log10 p-value. A −log10 p-value of 1.3 is equivalent to a permuted p-value of 0.05 (*p*<0.05*). Synonymous (BLACK) and non-synonymous SNPs (RED) are indicated. Regions informative for ocular localization and genes of interest are labelled in BLUE.

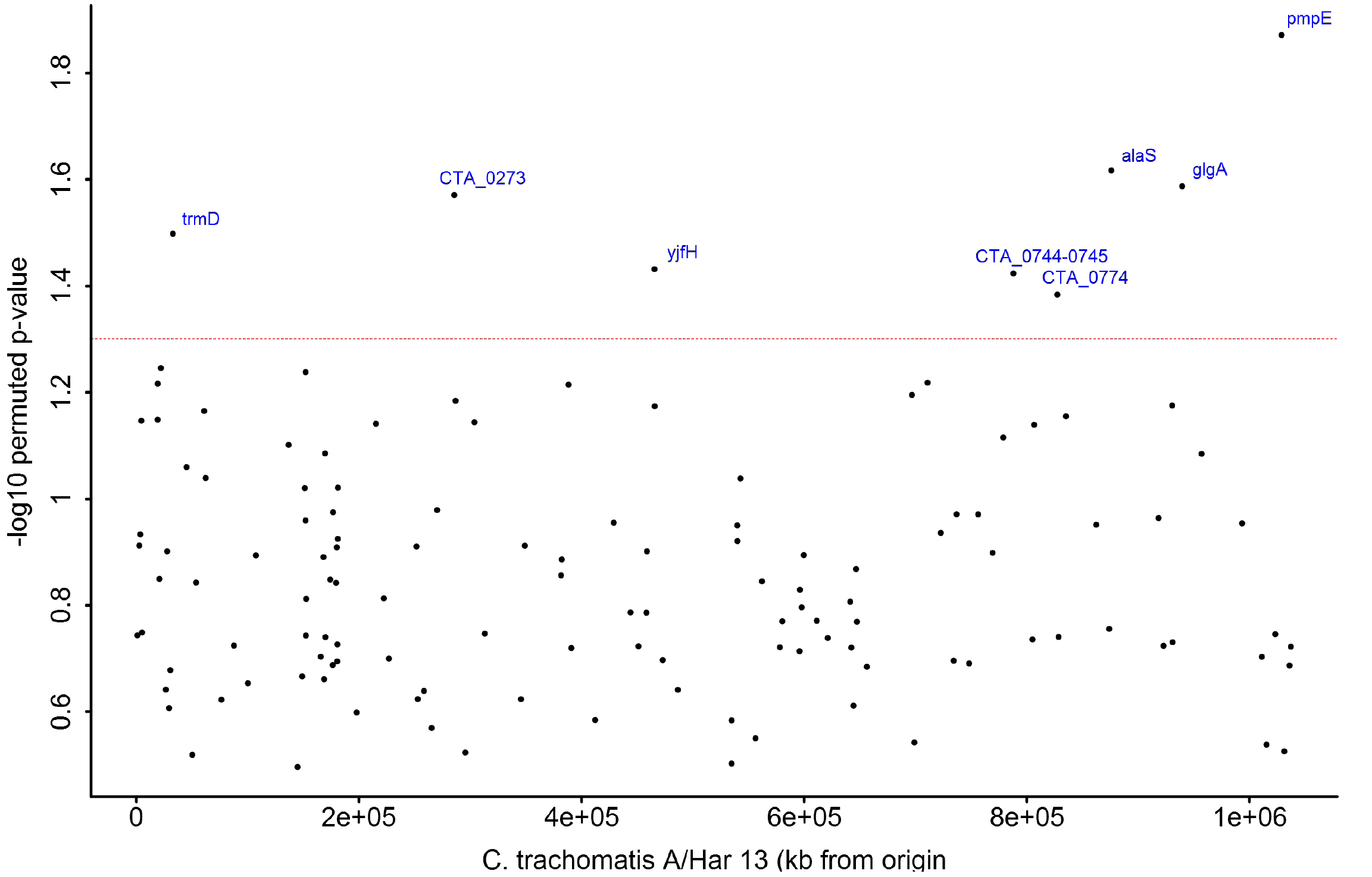
(B) Disease severity-associated SNPs across the *Ct* genome. From 129 SNPs identified in coding and non-coding regions, SNPs associated with the disease severity phenotype at genome-wide significance are identified using permutation-based ordinal logistic regression models adjusting for age in the *Ct* genome-wide association analysis. The threshold for genome-wide significance is indicated by the dashed line (*p*<0.05*). The y-axis shows the −log10 p-value. A log10 p-value of 1.3 is equivalent to a permuted p-value of 0.05 (*p*<0.05*). Genes significantly associated with disease severity are labelled in BLUE.

### Markers of disease severity in ocular C. trachomatis infection

Using permutation-based resampling methods, eight SNPs were found to be significantly associated with disease severity (*Figure 4(b)*). Seven of these are in coding regions (relative to *Ct A/HAR-13*). Five are present at nucleotide positions 465330 (OR=0.13, *p*=0.037*), 32779 (OR=0.12, *p*=0.032*), 875804 (OR=0.10, *p*=0.024*), 939488 (OR=0.10, *p*=0.026*) and 1028728 (OR=0.08, *p*=0.013*) (where p* is the permuted p-value with a genome-wide threshold of 0.05) representing synonymous codon changes within the genes *yjfH*, *trmD*, *alaS, glgA* and *pmpE* respectively. Three further genome-wide significant synonymous SNPs were present at positions 827184 (OR=0.3, *p*=0.041*) within the predicted coding sequence (CDS) *CTA0744*, 285610 (OR=0.12, *p*=0.027*) within *CTA0273* and 787841 (OR=0.13, *p*=0.043*) in the intergenic region between loci *CTA0744-CTA0745* (*Table 2(b)* and *Supplementary Information S6*).

## DISCUSSION

This is the first collection of clinical ocular *Ct* WGS from a single trachoma-endemic population to be characterized, enabling us to describe the population diversity of naturally occurring *Ct* in a treatment-naïve population. We used detailed clinical grading combined with microbial quantitation to perform a genome-wide association study (GWAS) and investigated associations between *Ct* polymorphisms with ocular localization and disease severity in trachoma.

Unlike the recently published Australian *Ct* sequences^10^, all Bijagós sequences clustered as expected within the T2 ocular clade derived from an urogenital ancestor^46,52^, each with loci typically associated with ocular tissue localization (*trpA* and *tarP*). Although the Bijagós sequences conform to the classical ocular genotype, the phylogenetic data show greater than expected diversity compared to historical reference strains of ocular *Ct*^4^ and a population of clinical ocular *Ct* sequences obtained from cultured clinical conjunctival swab specimens collected from another African trachoma-endemic population^53^ (*Supplementary Information S7*). Our use of direct WGS from clinical samples reveals the natural diversity of a population-based collection of endemic treatment-naïve ocular *Ct* infections. This diversity may indicate genome-wide selection for advantageous mutations as demonstrated in other pathogens^54^ or evidence that these are remnants of a previously larger and more diverse population in West Africa.

The apparent village-level clustering provides new evidence that WGS has the necessary molecular resolution to fully investigate *Ct* transmission. Although the number of sequences from each village were very small, overall *Ct* genomic diversity supports our hypothesis of ongoing or recent transmission, since diversity requires mutation, recombination and gene flow. The data from this study demonstrates such mutation and indicates that WGS data may be useful in defining transmission networks and developing transmission maps, which have not been adequately defined using alternative *Ct* genotyping systems. Whole genome mapping has previously been shown to be a useful tool in the analysis of outbreaks and bacterial pathogen transmission^55,56^ and thus has multiple potential applications in epidemiological analysis and transmission studies. However, greater numbers of sequences per village are required to validate this finding.

Such diversity is likely to be representative of recombination present in *Ct*^57^. Genome-wide recombination was common and widespread within these sequences. Extensive recombination has been noted in previous studies, and is thought to be a source of diversification with possible interstrain recombination^4,57^. Recombination may represent fixation of recombination in regions that are under diversifying selection pressure^4^.

Recently a handful of bacterial GWAS studies have provided insight into the genetic basis of bacterial host preference, antibiotic resistance, and virulence^58–63^. Until now, most inferences regarding disease-modifying virulence factors in chlamydial infection have been derived from a limited number of comparative genomic studies where only a few virulence factors were associated with disease severity. Chlamydial genomic association data have previously been used to highlight genes potentially involved in pathoadaptation^10,64^ and tissue localization^65^.

In the current GWAS we found 21 genome-wide significant non-synonymous SNPs associated with ocular localization and eight genome-wide significant synonymous SNPs associated with disease severity.

Confidence that new SNPs identified in the ocular localization GWAS are candidate markers of pathoadaptation is supported by the observation that half of the SNPs identified have previously been described as polymorphic or recombinant within *Ct* and the ocular serovars^8,66–68^. Genes expressed early in the *Ct* developmental cycle (with a peak at six hours post-infection [HPI]) or very late in the *Ct* developmental cycle (with a peak after 24 HPI) were over-represented, supporting the hypothesis that early events in infection and intracellular growth are crucial events in *Ct* survival and pathogenicity. Amongst the early-expressed genes are *CTA0156* (encoding early endosomal antigen 1 [EEA1]^69^), *CTA0498* (encoding translocated actin-recruiting phosphoprotein [*tarP*]^70^) and *CTA0884* (encoding polymorphic membrane protein D [PmpD]^71^), which have identified roles in entry to and initial interactions with host cells.

The eight disease severity-associated SNPs are within less well-characterized genes, although there is some evidence of correlation of expression in the *Ct* developmental cycle (with a peak between 18 and 24 HPI). Apart from *pmpE*, there is a paucity of published data showing polymorphism in these genes. This suggests that these SNPs may be important in ocular *Ct* pathogenesis, rather than in longer-term chlamydial evolution. Three of these genes are putative *Ct* virulence factors, with functions in nutrient acquisition (*glgA*^24,28,72^), host-cell adhesion (*pmpE*^73^) and response to IFNγ-induced stress (*trmD*^69^). Homologues of *alaS*^74,75^ and *CTA0273*^76,77^ are known virulence factors in related Gram-negative bacteria, suggesting that these genes are potentially important in *Ct* pathogenesis.

Transcriptome analysis of chlamydial growth *in vitro* has shown that there is highly upregulated gene expression of *trmD* (encoding a tRNA methyl–transferase) associated with growth in the presence of interferon gamma, thought to be important in the maintenance of chlamydial infection^69^. *yjfH* (renamed *rlmB*) is phylogenetically related to the TrmD family and encodes the protein RlmB, which is important for the synthesis and assembly of the components of the ribosome^78^. In *Escherichia coli*, *Haemophilus influenzae* and *Mycoplasma genitalium*, RlmB catalyses the methylation of guanosine 2251 in 23S rRNA, which is of importance in peptidyl tRNA recognition but is not essential for bacterial growth^78,79^. *alaS* encodes a tRNA ligase of the class II aminoacyl-tRNA synthetase family involved in cytoplasmic protein biosynthesis. It is not known to have virulence associations in chlamydial infection, but has been described as a component of a virulence operon in *Haemophilus ducreyi*^74^ and *H. influenzae*^75^. The CDS *CTA0273* encodes a predicted inner membrane protein translocase component of the autotransporter YidC, an inner membrane insertase important in virulence in *E. coli*^76^ and *Streptococcus mutans*^77^. This is the first study suggesting that these loci may be important in disease severity and host-pathogen interactions in chlamydial infection. A summary of available literature for these key ocular localization and disease severity-associated SNPs is tabulated in *Supplementary Information S8*. We cannot speculate further on the effect polymorphisms on expression. It is possible that the synonymous disease severity-associated SNPs are linkage-markers for disease-causing alleles that were not included in the GWAS. For both analyses, further mechanistic studies are required to establish causality, validity and to fully understand the nature of the associations presented in this study.

Though we were intrinsically limited to those cases where infection was detectable and from which we were able to obtain *Ct* WGS data, our population-based treatment naïve sample attempts to provide a representative picture of what is observed in ocular *Ct* infection. We acknowledge that there may be *Ct* genotypes that are cleared by the immune system such that we do not capture them a cross-sectional study. We are limited to the small sample size in this study, but attempt to address the issues of statistical power and multiple testing by using a bi-dimensional conjunctival phenotype and permutation-based multivariable regression analysis. To date the majority of published microbial GWAS have sample sizes under 500^80^, including several key studies examining virulence^59^ and drug-resistance^60^ in *Staphylococcus aureus* with sample sizes of 75 and 90 respectively.

The potential of bacterial GWAS has only recently been realized, and despite the limitations with sample size, its use to study *Ct* in this way is particularly important, since *in vitro* models are intrinsically difficult to develop and it has not been possible to study urogenital *Ct* in the same way due to the lack of a clearly defined *in vivo* disease phenotype. The genomic markers identified in this study provide important direction for validation through *in vitro* functional studies and a unique opportunity to understand host-pathogen interactions likely to be important in *Ct* pathogenesis in humans. The greater than expected diversity within this population of naturally circulating ocular *Ct* and the clustering at village-level demonstrates the potential utility of WGS in epidemiological and clinical studies. This will enable us to understand transmission in both ocular and urogenital *Ct* infection and will have significant public health implications in preventing and eliminating chlamydial disease in humans.

## METHODS

### Survey, Clinical Examination and Sample collection

Survey, clinical examination and sample collection methods have been described previously^45,81^. Briefly, we conducted a cross-sectional population-based survey in trachoma-endemic communities on the Bijagós Archipelago of Guinea Bissau. The upper tarsal conjunctivae of each consenting participant were examined, digital photographs were taken, a clinical trachoma grade was assigned and two sequential conjunctival swabs were obtained from the left upper tarsal conjunctiva of each individual using a standardized method^45^. DNA was extracted and *Ct omcB* (genomic) copies/swab quantified from the second conjunctival swab using droplet digital PCR (ddPCR)^81,82^.

We used the modified FPC (Follicles, Papillary hypertrophy, Conjunctival scarring) grading system for trachoma^39^. The modified FPC system allows detailed scoring of the conjunctiva for the presence of follicles (F score), papillary hypertrophy (conjunctival inflammation) (P score) and conjunctival scarring (C score), receiving a grade of 0-3 for each parameter. A single validated grader conducted the examinations and these were verified by an expert grader (masked to the field grades and ddPCR results) using the digital photographs. Grader concordance was measured using Cohen’s Kappa, where a Kappa > 0.9 was used as the threshold to indicate good agreement.

Conjunctival inflammation (P score) is known to have a strong association with *Ct* bacterial load in this and other populations^35,83–86^. For this study we used principal component analysis (PCA) to combine the presence of inflammation (defined by the P score using the FPC trachoma grading system^39^) with *Ct* bacterial load (defined by tertile cut-offs illustrated in *Supplementary Information S9*)^87^. The conjunctival disease phenotype is a dimension reduction of these two variables defining what we observed in the conjunctiva at the time of sampling (*Figure 5*). Dimension reduction using PCA to define complex disease phenotypes in GWAS is well-recognised, as it allows multiple traits to be included to capture a more complex phenotype and accounts for correlation between traits. This approach therefore may reveal novel loci or pathways that would not be evident in single-trait GWAS, where the full extent of genetic variation cannot be captured^40^.

**Figure 5.**
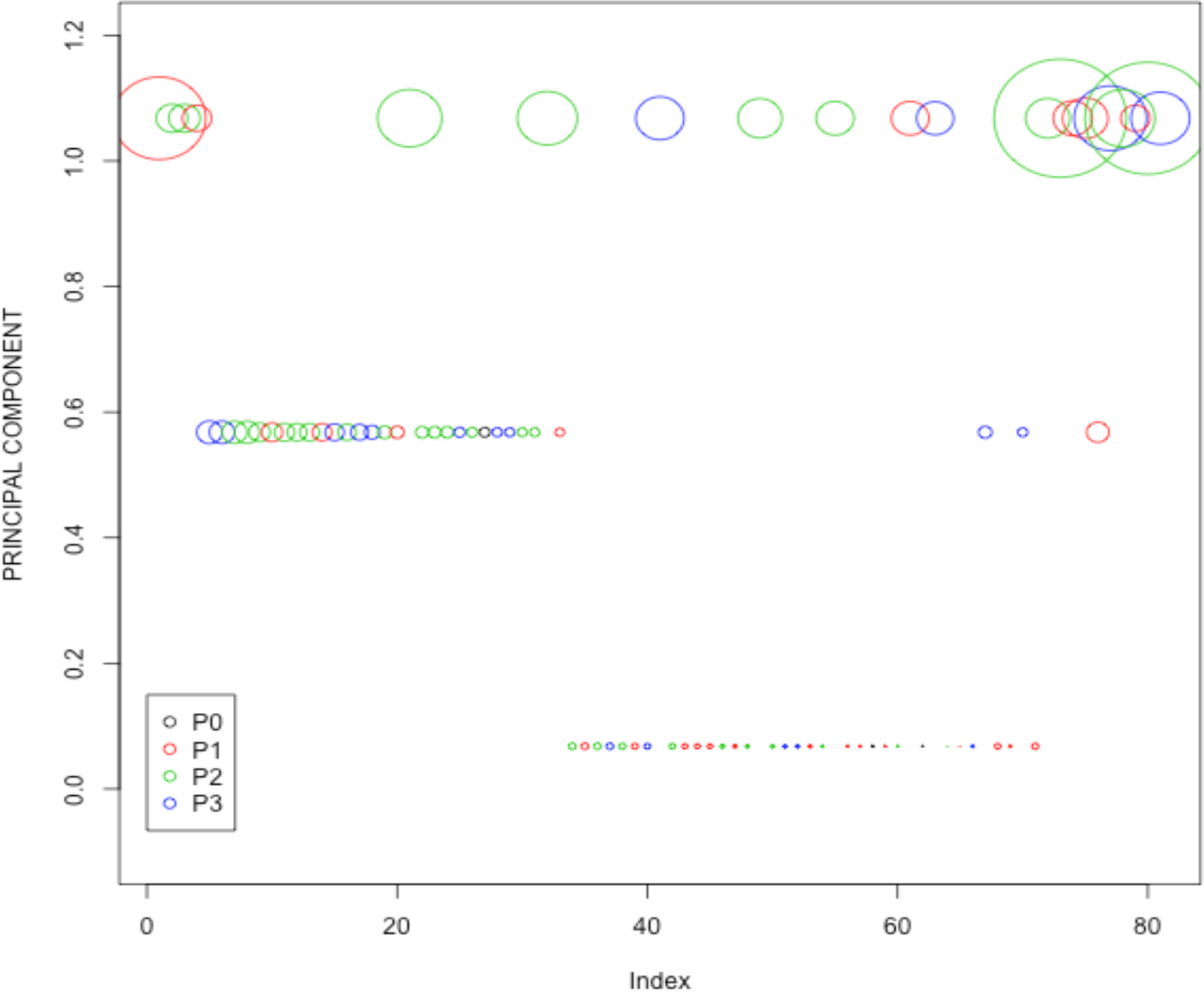
Composite *in vivo* conjunctival disease severity phenotype in ocular *Chlamydia trachomatis* infection. A composite *in vivo* phenotype was derived using principal component analysis (PCA) for dimension reduction of two phenotypic traits: a disease severity score (using the P score value) and *C. trachomatis* load (where *C. trachomatis* load was log transformed and cut-offs determined from the resulting density plot (See *Supplementary Information S9*)). Each circle represents an individual infection (represented on the x axis (Index), n=81). Circle size reflects *C. trachomatis* load and circle colour reflects inflammatory P score (P0-P3) defined using the modified FPC (Follicles, Papillary Hypertrophy, Conjunctival Scarring) grading system for trachoma^39^

### Preparation of chlamydial DNA for cell culture

For eight specimens, whole genome sequence (WGS) data were obtained following *Ct* isolation in cell culture (from the first conjunctival swab). Briefly, samples were isolated in McCoy cell culture by removing 100μl eluate from the original swab with direct inoculation onto a glass coverslip within a bijou containing Dulbecco’s modified Eagles’ Medium (DMEM). The inocula were centrifuged onto cell cultures at 1800rpm for 30 minutes. Following centrifugation the cell culture supernatant was removed and cycloheximide-containing DMEM added to infected cells which were then incubated at 37°C in 5% CO_2_ for three days. Viable *Ct* elementary bodies (EB) were observed by phase contrast microscopy. Cells were harvested and further passaged every three days until all isolates reached a multiplicity of infection between 50-90% in 2xT25 flasks. Each isolate was prepared and EBs purified as described previously^88^. DNA was extracted from purified EBs using the Promega Wizard Genomic Purification kit according to the manufacturer’s protocol^89^.

### Pre-sequencing target enrichment

For the remaining specimens (*n=118*), WGS data were obtained directly from clinical samples. DNA baits spanning the length of the *Ct* genome were compiled by SureDesign and synthesized by SureSelect^XT^ (Agilent Technologies, UK). *Ct* DNA extracted from clinical samples was quantified and carrier human genomic DNA added to obtain a total of 3μg input for library preparation. DNA was sheared using a Covaris E210 acoustic focusing unit^31^. End-repair, non-templated addition of 3’–A adapter ligation, hybridisation, enrichment PCR and all post- reaction clean-up steps were performed according to the SureSelect^XT^ Illumina Paired-End Sequencing Library protocol (V1.4.1 Sept 2012). All recommended quality control measures were performed between steps.

### Whole genome sequencing and sequence quality filtering

DNA was sequenced at the Wellcome Trust Sanger Institute using Illumina paired-end technology (Illumina GAII or HiSeq 2000). All 126 sequences passed standard FastQC quality control criteria^90^. Sequences were aligned to the most closely related reference genome, *Chlamydia trachomatis A/HAR-13* (GenBank Accession Number NC_007429.1 and plasmid GenBank Accession Number NC_007430.1), using BWA^91^. SAMtools/BCFtools (SAMtools v1.3.1)^92^ and GATK^93^ were used to call SNPs. We used standard GATK SNP calling algorithms, where >10x depth of coverage is routinely used as the threshold value^93,94^. This has been shown to be adequate for SNP calling in this context^43,44,46,94^.

Variants were selected as the intersection dataset between those obtained using both SNP callers and SNPs were further quality-filtered. SNP alleles were called using an alternative coverage-based approach where a missing call was assigned to a site if the total coverage was less than 20x depth or where one of the four nucleotides accounted for at least 80% total coverage^95^. There was a clear relationship between the mean depth of coverage and proportion of missing calls, based on which we retained sequences with greater than 10x mean depth of coverage over the whole genome (81 sequences retained).

Heterozygous calls were removed and SNPs with a minor allele frequency of less than 25% were removed. Samples with greater than 25% genome-wide missing data and 30% missing data per SNP were excluded from the analysis (n=10, 71 sequences retained). The quality assessment and filtering process is shown in *Figure 1*. Detail of WGS data is contained in *Supplementary Information S10*.

### Phylogenetic Reconstruction

Samples were mapped to the ocular reference strain *Ct A/HAR-13* and SNPs were called as described above. Phylogenies were computed using RAxML version 7.8.2^96^ from a variable sites alignment using a GTR+gamma model and are midpoint rooted. Recombination is known to occur in *Ct*^4,6^ and can be problematic in constructing phylogeny. We applied three compatibility-based recombination detection methods to detect regions of recombination using PhiPack^97^: the pairwise homoplasy index (Phi), the maximum Chi2 and the neighbour similarity score (NSS) across the genome alignment. We also examined the confidence in the phylogenetic tree by computing RAxML site-based likelihood scores^96^. Phylogenetic trees were examined adjusting for recombination using the methods described above.

Additionally, sequence data for the tryptophan operon (*CTA0182* and *CTA0184-CTA0186*), *tarP* (*CTA0498*), nine polymorphic membrane proteins (*CTA0447-CTA0449, CTA0884, CTA0949-CTA0952* and *CTA0954*) and *ompA* (*CTA0742*) were extracted from the 81 ocular *Ct* sequences from Guinea Bissau retained after quality control filtering described above, 48 ocular sequences originating from a study conducted in Kahe village, Rombo District, Tanzania^53^ and 38 publicly available reference sequences. Phylogenies were constructed as described above.

Polymorphisms, insertions and deletions (INDELs) and truncations for the tryptophan operon were manually determined from aligned sequences using SeaView^98^. Tyrosine repeat regions and actin-binding domains in *tarP* were found using RADAR^99^ and Pfam^100^ respectively.

### Pairwise diversity

A comparison was made between the two population-based *Ct* sequence data sets from the Bijagós (Guinea Bissau) and Rombo (Tanzania) sequences whereby short read data from the 81 Bijagós sequences and 48 Rombo sequences were mapped against *Ct A/HAR-13* using SAMtools. Within population pairwise nucleotide diversity was calculated using the formula:

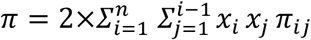

where n is the number of sequences, x is the frequency of sequences i and j and πij is the number of nucleotide differences per site between sequences i and j^101^. Frequency of sequences was considered uniform within the populations and sites with missing calls were excluded on a per sequence basis.

### Genome-Wide Association Analyses

To investigate the association between *Ct* polymorphisms with ocular localization and clinical disease severity, we used permutation-based logistic regression methods, which are powerful and well-recognised tools in GWAS, allowing for adjustment for population structure, age and gender in the model and accounting for multiple testing ^48–51^.

We used permutation analyses of 100,024 phenotypic re-samplings, where the distribution of the p-value was approximated by simulating data sets through randomisation under the null hypothesis of no association between phenotype and genotype. Genome-wide significance was determined as *p*≤0.05*, where *p\** was defined as the fraction of re-sampled (simulated) data that returned p-values that were less than or equal to the p-values observed in the data^87^. All analyses were conducted using the R statistical package v3.0.2 (The R Foundation for Statistical Computing, http://www.r-project.org) using MASS, GLM and lsr. All R script used for these analyses is contained within *Supplementary Information S11* and is released as a CC-BY open resource (CC-BY-SA 3.0).

#### Ocular localization

Tissue localization is defined as the localization (or presence) of a detectable *Ct* infection to either the conjunctival epithelium or the urogenital tract. Short read data from the 129 clinical ocular sequences from the pairwise diversity analysis and 38 publicly available reference sequences from ocular (n=8), urogenital (n=17) and rectal (n=13) sites were mapped against *Ct A/HAR-13* using SAMtools. Only polymorphic sites were retained and SNPs were filtered as described above. The final analysis includes 1007 SNPs from 157 sequences, a phylogeny of which is contained within *Supplementary Information S7*. A permutation-based generalized linear regression model was used to test the association between collection site (ocular or urogenital tissue localization) and polymorphic sites. For each SNP the standard error for the t statistic was estimated from the model and used to calculate the odds ratios (OR) and 95% confidence intervals. A χ^2^ test was used to determine the association between ocular localization-associated SNPs and both gene expression stage and predicted localization of the encoded proteins. Developmental cycle expression stage for each transcript was based on data and groupings from Belland *et al.*^69^. Predicted localization of expressed proteins was defined using the consensus from three predictions using Cello^102^, pSORTB^103^ and LocTree3^104^.

#### Clinical Disease Severity

A permutation-based ordinal logistic regression model was used to test the association between the disease severity score (using the *in vivo* conjunctival phenotype defined previously) and polymorphic sites. The final analysis includes 129 SNPs from 71 sequences derived as described in *Figure 1*. For each SNP the standard error for the t-statistic was estimated from the model and used to calculate the odds ratios (OR) and 95% confidence intervals. Individuals’ age and gender were included as a covariate to the regression analysis.

We investigated the effect of population structure on the results of the GWAS analysis using PCA^105^. The first three principal components (PC) captured the majority of structural variation but including these in the model had no effect and therefore these were not included in the final model.

We corrected for genomic inflation if the occurrence of a polymorphism in the population was over 90% or there was a minor allele frequency of 3%.

## DATA AVAILABILITY

All sequence data is available from the European Bioinformatics Institute (EBI) short read archive. See *Supplementary Information S12* for details and accession numbers.

## ACKNOWLEDGEMENTS

AL was funded by the Wellcome Trust through a Clinical Research Training Fellowship (grant number 097330/Z/11/Z). MH and ChR were funded by a Wellcome Trust Program Grant (grant number 079246/Z/06/Z). ChR was funded by the Wellcome Trust Institutional Strategic Support Fund (grant number 105609/Z/14/Z). Work undertaken by the Wellcome Trust Sanger Institute by NRT, HSS and JH was supported by the Wellcome Trust (grant number 098051 and the Wellcome Trust Institutional Strategic Support Fund (grant number 105609/Z/14/Z)). TGC is funded by the Medical Research Council UK (grant numbers MR/K000551/1, MR/M01360X/1, MR/N010469/1). JP is funded by a BBSRC PhD studentship, and FC and HP were funded by Bloomsbury Colleges Research Fund PhD studentships. We extend thanks to colleagues at the Programa Nacional de Saúde de Visão in the Ministério de Saúde Publica in Bissau, the study participants and dedicated field research team in Guinea Bissau and the MRC Unit The Gambia for their support and collaboration in this work. All communities received treatment for endemic trachoma in accordance with WHO and national policies following the survey.

## FOOTNOTES

ARL/RLB/MJH designed the study. ARL/SEB/EC/MN conducted the field study. ARL/ChR/SEB conducted the molecular laboratory work. LTC/IN performed the Chlamydial cell culture. HSS/JH designed and performed the whole genome sequencing and QC. ARL/ChR/HP/FC/TC conducted the GWAS analysis. HP/JP/SH/JH/HSS supported the phylogenetic analysis. ARL/HP/MJH/DCWM/TC/NRT wrote the paper. All authors have contributed to and reviewed the manuscript.

